# Fluid intelligence and naturalistic task impairments after focal brain lesions

**DOI:** 10.1101/2021.05.12.443802

**Authors:** Verity Smith, Clara Pinasco, Jascha Achterberg, Daniel J. Mitchell, Tilak Das, Maria Roca, John Duncan

**Author notes:** These authors contributed equally to the work. Correspondence, MRC Cognition and Brain Sciences Unit, CB2 7EF, UK.

## Abstract

Classical executive tasks, such as Wisconsin card-sorting and verbal fluency, are widely used as tests of frontal lobe control functions. Since the pioneering work of Shallice and Burgess (1991), it has been known that complex, naturalistic tasks can capture deficits that are missed in these classical tests. Matching this finding, deficits in several classical tasks are predicted by loss of fluid intelligence, linked to damage in a specific cortical “multiple-demand” (MD) network, while deficits in a more naturalistic task are not. To expand on these previous results, we examined the effect of focal brain lesions on three new tests – a modification of the previously-used Hotel task, a new test of task switching after extended delays, and a test of decision-making in imagined real-life scenarios. As potential predictors of impairment we measured volume of damage to *a priori* MD and default mode (DMN) networks, as well as cortical damage outside these networks. Deficits in the three new tasks were substantial, but were not explained by loss of fluid intelligence, or by volume of damage to either MD or DMN networks. Instead, deficits were associated with diverse lesions, and not strongly correlated with one another. The results confirm that naturalistic tasks capture cognitive deficits beyond those measured by fluid intelligence. We suggest, however, that these deficits may not arise from specific control operations required by complex behaviour. Instead, like everyday activities, complex tasks combine a rich variety of interacting cognitive components, bringing many opportunities for processing to be disturbed.

---

It has long been thought that complex, open-ended tasks may capture aspects of frontal “executive” impairment that are missed in the more constrained setting of conventional neuropsychological testing. In a pioneering study, Shallice and Burgess (1991) introduced two tasks designed to mimic the open-ended character of everyday problem solving. In the 6-element task, the patient had to divide a period of 15 minutes between six different tasks, with freedom to switch tasks whenever they chose, but with several additional rules concerning task order and time allocation. In the multiple-errands task, the patient undertook a list of activities in a street of shops, again organizing the entire performance to respect a list of rules and requirements. In three frontal patients, Shallice and Burgess (1991) showed major impairment in these tasks, despite generally good performance on a battery of more conventional executive tests such as Wisconsin card sorting (Milner 1963), verbal fluency (Benton 1968) and Trails B (Reitan 1955).

In previous studies, we have investigated the link between executive tests and fluid intelligence, measured with a standard test such as the Culture Fair (Institute for Personality and Ability Testing 1973). For a number of conventional tests, including card sorting, fluency, and Trails, deficits in several patient groups are largely explained by a loss of fluid intelligence; once fluid intelligence is partialled out, performance is largely equivalent for patients and controls (Roca et al. 2010, 2012, 2013, 2014). Fluid intelligence deficits are linked to damage in a distributed cortical “multiple-demand” or MD network, incorporating specific regions of lateral frontal, dorsomedial frontal, insular and parietal cortex (Woolgar et al. 2010, 2018; Barbey et al. 2012); for evidence on white matter connections (see Gläscher et al. 2010). Performance on tests such as card sorting, fluency, and Trails may largely reflect the functions of this network. Findings are different for a more open-ended task, the Hotel, designed to mimic the Shallice and Burgess (1991) 6-element task in a more realistic setting (Manly et al. 2002). For the Hotel task, we have repeatedly found that performance is only weakly related to fluid intelligence, and partialling fluid intelligence does not remove patient deficits (Roca et al. 2010, 2012, 2013, 2014). These findings suggest less specific dependence on MD functions.

In the present work, we extended these prior findings using three new tests, administered along with the Culture Fair to a group of patients with focal lesions in different regions of cortex. First, we used a version of the Hotel, somewhat shortened compared with previous versions. Second, we designed a new test of everyday comprehension and problem-solving, based on short vignettes describing real-life situations and their accompanying decisions. Third, we designed a new task-switching test to mimic just one aspect of the complex processing requirements of the Hotel. It has often been suggested that, in this test, patients may fail to divide their time between component sub-tasks because, having become immersed in one, they forget the larger requirement to give some time to all of them (Manly et al. 2002). To investigate this kind of immersion as a possible key factor in naturalistic behaviour, we modified a standard task-switching paradigm (Rogers and Monsell 1995) to manipulate the length of time before a task switch. Compared to our other two tasks, this one focused on a quite specific cognitive requirement, but one that we thought might be important in temporally extended, open-ended behaviour.

As predictors of cognitive impairment, we considered damage to two *a priori* networks. The first was the MD network, predicted to be important for fluid intelligence but less so for the other tasks. The second was the default-mode network (DMN). In functional brain imaging, the DMN is well known as a set of brain regions with strong functional connectivity (Yeo et al. 2011), deactivation in many tasks compared to rest (Shulman et al. 1997), but positive activation linked to mind-wandering (Mason et al. 2007; Christoff et al. 2009), self-related thought (D’Argembeau et al. 2005), and sometimes large externally-directed task switches (Crittenden et al. 2015; Smith et al. 2018). As regards the Hotel, one specific reason for suspecting DMN involvement came from the previous results of Roca et al. (2010, 2011), who linked Hotel deficits to anterior frontal lesions. On the medial surface, anterior frontal cortex includes a large region of the DMN, and though Roca et al. (2011) combined lateral and medial patients, it seemed possible that the DMN component was responsible for Hotel deficits. More broadly, a large body of imaging work links the DMN to both social cognition and imagination of cognitive scenes or episodes (Frith and Frith 2003; Amodio and Frith 2006; Addis et al. 2007, 2009; Andrews-Hanna et al. 2010). Several authors have suggested that situation representations in the DMN place ongoing cognition in a broader context (Hassabis and Maguire 2007; Ranganath and Ritchey 2012; Smith et al. 2018). Reference to a larger context could be especially important in naturalistic problem-solving, including behavioural management over an extended time period. For each patient, we measured volume of damage in MD, DMN and other cortical regions, and linked these lesion measures to behavioural impairment.

A different possibility is that naturalistic task deficits do not reflect specific control requirements associated with complex behaviour. Rich, varied tasks based on the requirements of everyday cognition are likely by definition to have many different cognitive components, dependent on multiple brain processes (Burgess et al. 2000). These components, furthermore, are unlikely to act independently; if one component is weakened through brain damage, it may send delayed or inaccurate input to others, or compete for mental resources. Of course, all tasks are potentially influenced by multiple sensory, motor and cognitive functions, but for rich tasks, it may be especially unlikely that one or a few core processes are responsible for the bulk of neuropsychological impairments. On this account, complex tasks are highly sensitive to brain damage simply because there are many opportunities for their processing to be disturbed. The prediction would be deficits that are not well explained by any one focal cognitive impairment, or by damage to any one cortical region or network.

## Methods

### Participants

A convenience sample of 34 patients was recruited from the Cambridge Cognitive Neuroscience Research Panel at the MRC Cognition and Brain Sciences Unit. The study was conducted in accordance with ethical approval from the Cambridge Psychology Research Ethics Committee. Patients were selected on the basis of having chronic, focal lesions from mixed aetiology excluding traumatic brain injury, and aged between 18-80 years old. There were no other formal exclusion criteria for region of lesion or specific cognitive deficit. Two patients were unable to complete more than 1 task and were excluded from further analysis. Demographic and lesion information for the remaining 32 patients is presented in Table 1. 30 non-brain-damaged control participants were also tested. To ensure good correspondence between patient and control groups on factors of age and years of educations, data for the patient group was collected first and control participants were then selected to approximate the patient distribution. Accordingly, patient and control groups did not differ significantly in age (patients mean =58.4 years, SD=15.3; controls mean=52.6, SD=18.9; t(61)=1.35, p=0.18) or years of education (patients mean=13.9, SD=2.3; controls mean=14.7, SD=3.9; t(61)=1.04, p=0.30).

**Table 1.** Clinical and demographic data. For some patients precise lesion date is unknown and an estimate is given based on date of joining panel.

| ID | Age (years) | Years of Education | Aetiology | Lesion hemisphere | Time between lesion and tests (years) |
| --- | --- | --- | --- | --- | --- |
| 1 | 73 | 11 | stroke | right | 11.5 |
| 2 | 32 | 16 | stroke | right | 8.5 |
| 3 | 47 | 11 | tumour | right | 1.5 |
| 4 | 74 | 11 | tumour | right | >15 |
| 5 | 71 | 11 | tumour | right | 9 |
| 8 | 53 | 16 | tumour | right | 4 |
| 9 | 70 | 16 | tumour | left | 6.5 |
| 10 | 71 | 17 | stroke | right | 17 |
| 11 | 67 | 16 | tumour | left | 25.5 |
| 12 | 79 | 9 | stroke | right | 20 |
| 13 | 36 | 14 | tumour | left | 7 |
| **14** | 54 | 16 | stroke | right | 18 |
| **15** | 71 | 16 | stroke | right | 20 |
| **16** | 78 | 13 | stroke | right | 14 |
| **17** | 62 | 13 | resection | right | 19 |
| **18** | 54 | 14 | abscess | left | 13 |
| **19** | 68 | 11 | tumour | bilateral | 20 |
| **20** | 53 | 11 | tumour | left | 19 |
| **21** | 35 | 13 | tumour | right | 7.5 |
| **22** | 65 | 14 | infection | left | 12.5 |
| **23** | 42 | 16 | tumour | left | 9 |
| **24** | 54 | 11 | tumour | right | 14 |
| **25** | 30 | 13 | tumour | left | 3 |
| **26** | 47 | 13 | tumour | right | 13 |
| **27** | 73 | 17 | tumour | left | 3 |
| **28** | 60 | 13 | tumour | left | 19 |
| **29** | 77 | 18 | tumour | right | 13.5 |
| **30** | 25 | 14 | tumour | right | 5 |
| **31** | 46 | 16 | tumour | bilateral | 13 |
| **32** | 67 | 13 | tumour | bilateral | 8 |
| **33** | 64 | 16 | tumour | left | 18 |
| **34** | 67 | 16 | unknown | right | >18 |

### Testing

Participants were given a battery of computer-based and other tasks described below. The test battery was completed in a single session lasting around 90-120 minutes. The battery consisted of 7 tasks, presented in fixed order. In this paper we present data for Culture Fair (presented first in the session), Situations (second), Hotel (fourth and seventh; see below), and Switch Time (sixth). The remaining three tests were included for other purposes, not concerned with naturalistic decision making. They were a more conventional task switching paradigm (presented third), adapted from Smith et al. (2018), an object in place concurrent discrimination memory task (presented fifth), similar to Gaffan (1994), and a comparison of attention control by scene or object cues (presented last). Four patients did not complete the Switch Time task due to fatigue. Additionally, for one of these four patients, the battery was split into two shorter sessions. Computer-based tasks were given on a Dell 1280×1024 resolution laptop, controlled using Psychophysics Toolbox for MATLAB (Brainard 1997).

### Tasks

#### Culture Fair

Participants were given the standardised version of the Cattell Culture Fair, Scale 2 Form A (Institute for Personality and Ability Testing 1973) consisting of four timed subtasks (series completion, odd one out, matrices, topological relations). At the start of each subtask, the experimenter read aloud the rules to the participant and went through 3 examples with them. Total correct scores were calculated and then converted into IQs from the standardised table of norms. To match the rest of the data set, IQ scores were inverted such that higher numbers would correspond to poorer performance.

#### Situations

The Situations task was designed to test social and non-social decision-making in real-life vignettes. During the task, participants were shown 12 short stories on the computer screen, and after each story asked one social judgement question, one emotion judgement question and one executive judgement question. An example story and question set is presented in Figure 1. The full set of stories and questions is presented in the Supplementary Materials. For each item, participants were first shown the story text and asked to read through the story. After reading, participants were asked to press a button. With the story text still present on the screen, the questions were then presented one by one, along with 3 possible answers each. The answer options were designed such that one was correct, one was very incorrect and the third was plausible but less optimal than the correct option. Participants were asked to respond as quickly as they could using buttons “1”, “2” or “3” on the keyboard, corresponding to the 3 possible answers. The order of story presentation was randomised and the order in which the questions were presented was pseudorandomised such that each type of question (social, emotional and executive) was presented equally often first, second and third. The position of the correct answer was counterbalanced across question types such that it appeared in positions 1, 2 and 3 on an equal number of trials.

**Figure 1.**
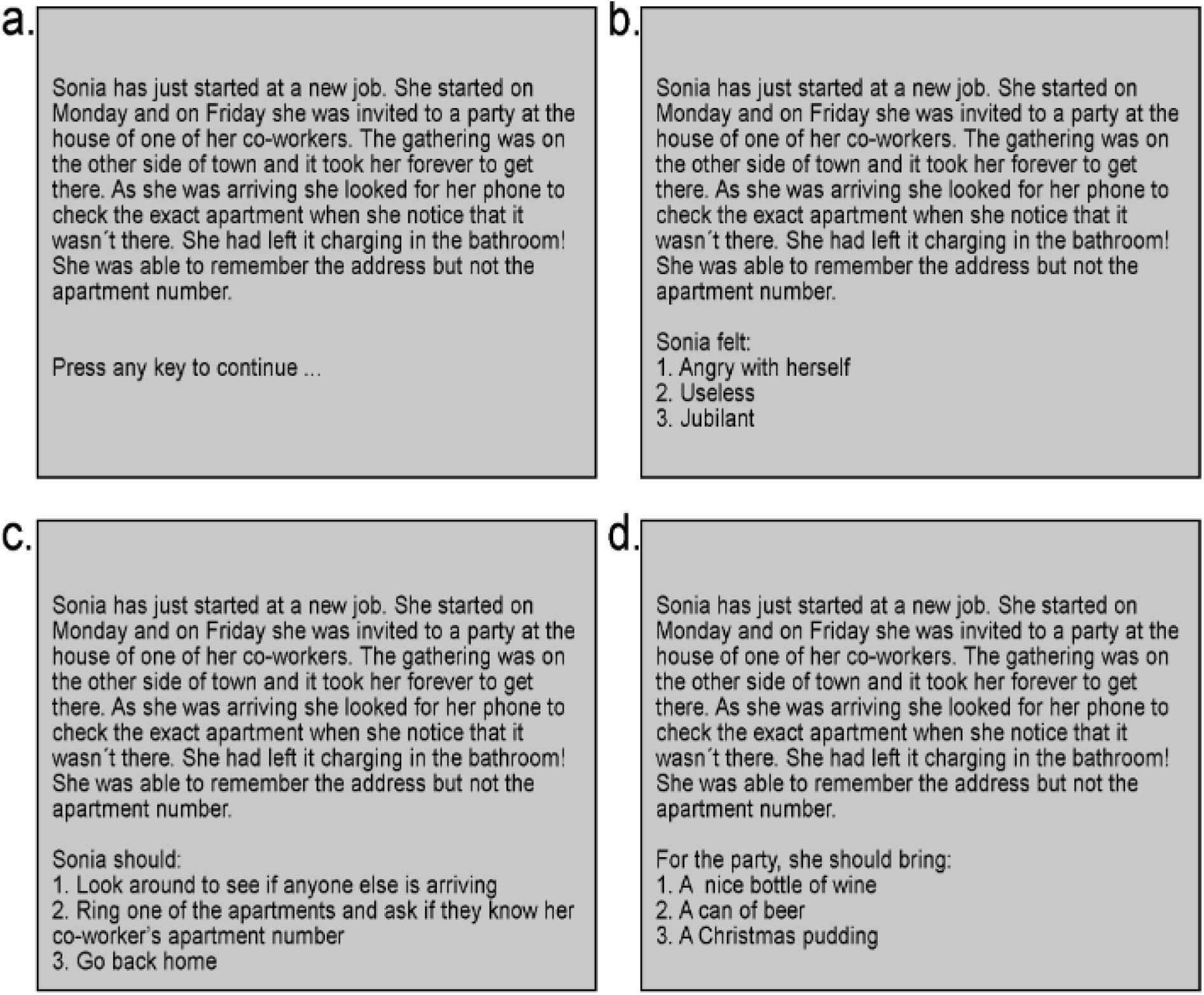
An example item in the Situations task. (a) Participants first read the story at their own pace before pressing any key to continue. Participants were then presented with an emotion question (here b), an executive question (here c) and a social question (here d), in counterbalanced order. The questions were self-paced and the story remained on the screen throughout the question phase.

We scored proportion error and median response time (RT) for correct trials. RTs for the three different question types were strongly correlated for both patients (for the three possible pairs of question types, r = .89 to .95) and controls (.49 to .71). Proportion error scores for the three question types were also correlated in patients (.22 to .59), though not in controls (-.03 to .11), perhaps in part because error proportions were low. As overall measures of performance, we averaged RTs and proportion error scores across the three question types.

#### Hotel

The Hotel task used materials laid out on a table in front of the participant. An example of the task layout is presented in Figure 2. Participants were asked to imagine they were in a job interview for a position at a hotel and were asked to try 3 different hotel activities, each one involving sorting a stack of sheets of paper. Participants were told that they would have 9 minutes to work on the three activities and that it would be impossible to finish any of them completely in the time limit. Instead, they should ensure that some time was allotted to each activity. To keep track of the time participants were given a clock. Throughout the task the clock was turned away from the participants, but participants could choose to check it at any time before returning it to its backward-facing position.

**Figure 2.**
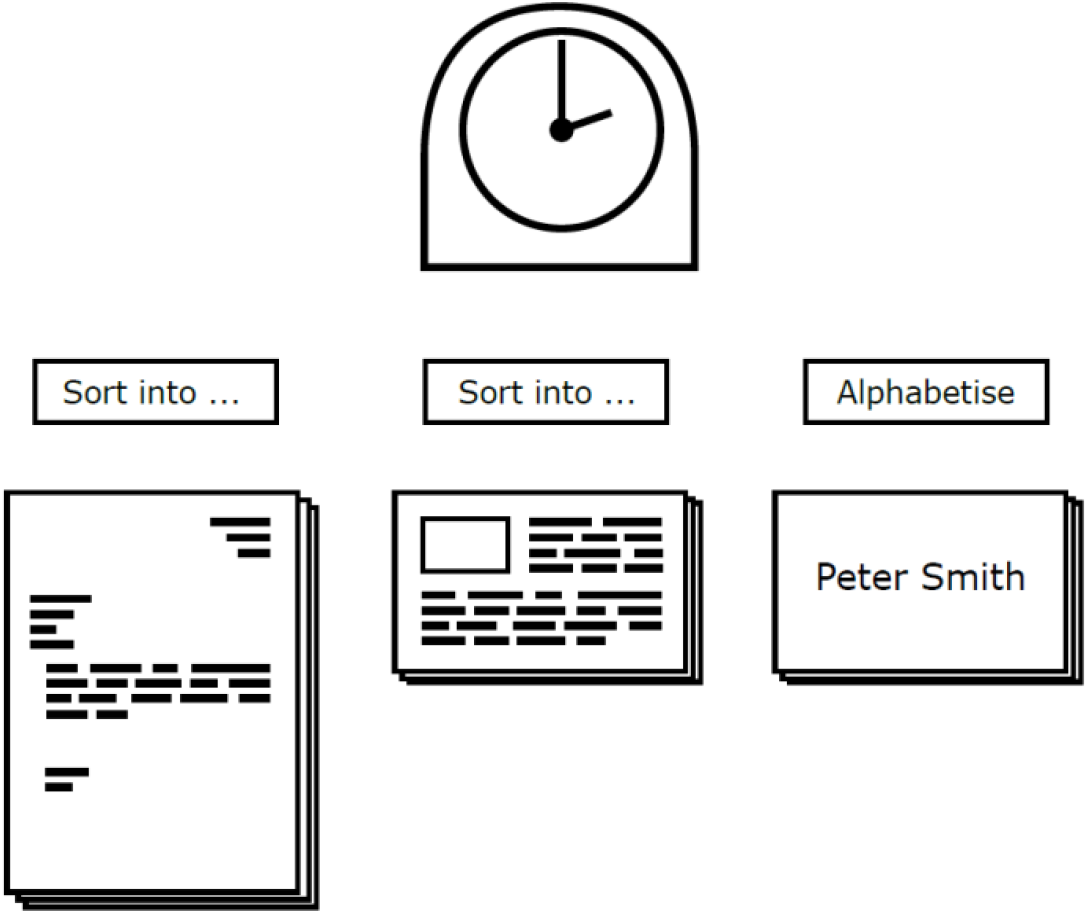
Schematic experimental set up for the Hotel task. Note that the clock was facing away from the participants except when explicitly checked.

Participants were given two variants of the task, which varied the form of periodic interruptions. The order of the two variants was counterbalanced across participants, with two other tasks performed between the two variants. Interruptions were motivated by the work of Manly et al. (2002), who found that performance improved when patients were given an occasional auditory alert designed to break focus on the current activity and reorient attention to the overall goal. In our version, interruptions were designed to reorient participants’ attention either within the current activity (internal interruption) or to the external environment (external interruption). In the internal interruption condition a yellow sheet of paper was placed after every 7 task items in each activity. Participants were told to place the yellow item in the same pile as the previous item. In the external interruption condition a written instruction, asking participants to perform an action directed towards an aspect of the surrounding environment, was placed after every 7 task items. Participants were asked to follow the written instruction (e.g. point to a window) and then place the instruction sheet to the side.

For the two task versions, there were two separate sets of 3 activities. The activity set paired with each interruption condition was counterbalanced across participants. Set A consisted of sorting conference name tags by alphabetical order, sorting invoices into piles according to the vendors, and sorting bills into piles according to customer name. Set B consisted of sorting staff name tags by alphabetical order, sorting restaurant lists into piles by their location, and sorting spa receipts into piles by treatment.

During the task, the experimenter kept a continuous record of which activity the participant was working on, using computer keys to indicate each time an interruption was encountered and each time the activity was switched, along with which new activity was begun. In line with previous work (Manly et al. 2002; Torralva et al. 2009; Roca et al. 2010), the score was the summed deviation from optimal time (180 seconds per activity) across the 3 activities. Preliminary analysis showed no differences between interruption types. However, there was evidence that participants (especially controls) improved their strategy when performing the task for the second time. Accordingly, scores were based just on the first version performed, whichever interruption type it involved.

#### Switch Time Task

One difference between the Hotel task and other classical switching tasks is the long period (∼180s) between switches. The Switch Time task was designed to test whether patients are particularly impaired at task switching after longer rather than shorter periods. As noted above, only 28 patients took part in this task. Task events are illustrated in Figure 3. For each trial, participants were presented with a picture to the left of the screen and a word with a letter missing on the right of the screen. Participants were required to make yes/no judgements on one of these stimuli based on a task rule cued by a central shape. If the rounded corners of the cue pointed towards the left, then participants were asked to do the picture task. If the rounded corners of the cue pointed towards the right, then participants were asked to do the word task. The two tasks used were taken from Crittenden et al. (2015). For pictures, the decision was whether the item would fit in a shoebox; for words, it was whether addition of a letter ‘a’ in the blank position would create a real word. Participants were asked to make a “yes” or “no” response, by left or right keypress respectively. Performance was self-paced, with the stimuli remaining until a key was pressed, and participants were asked to respond as quickly as possible without making mistakes. An inter-trial interval (ITI) of seconds followed each response.

**Figure 3.**
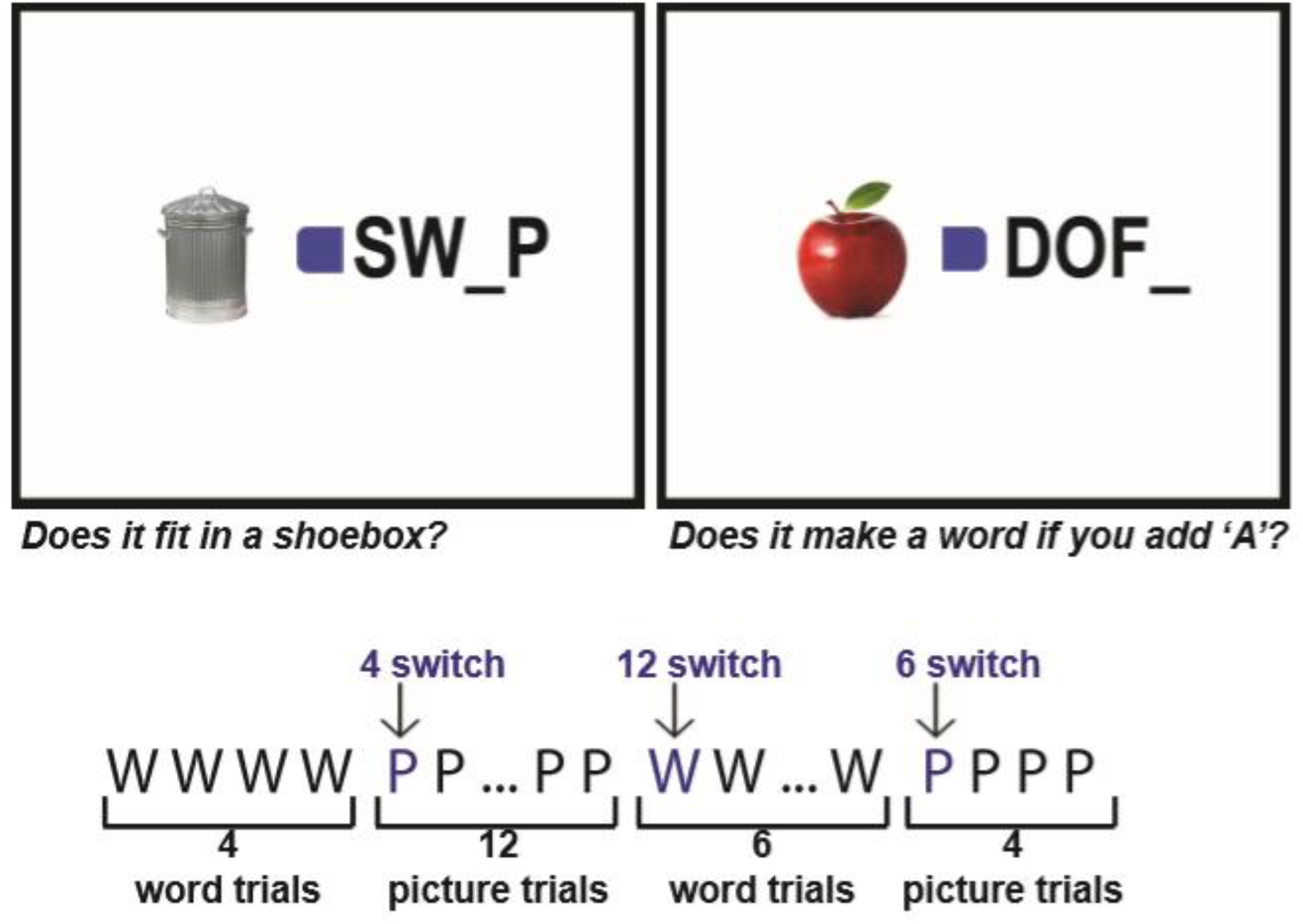
Task design for the Switch Time task. Pictures were presented to the left of the screen with words containing a blank space presented to the right of the screen. In the centre was a blue square with a curved edge made by removing upper and lower corners. The side of the curved edge instructed the participant to carry out the left hand picture task or the right hand word task. The cued side remained fixed for blocks of 4, 6 or 12 trials, creating switch trials which varied by number of trials of the previous task (switch 4, switch 6 and switch 12 trials). In the figure, task cues have been enlarged to increase visibility of curved edges.

Pilot studies revealed that switch performance systematically declined with increasing numbers of repeated trials prior to a cue change, reaching an asymptote at around 12 repeats. Accordingly, trials of the same task were repeated in blocks of 4, 6 or 12 trials, with block length presented in a pseudorandom order. Trials could thus be sorted into 4 transition types: stay trials (task trials preceded by the same task), switch 4 trials (task switch after 4 of the same task), switch 6 trials (task switch after 6 of the same task), and switch 12 trials (task switch after 12 of the same task). The experiment consisted of 1 run of 182 trials. The run contained 4 blocks of each length for each task, plus a final run of 6 trials after the last switch. A break was inserted into the run, approximately halfway through the trials, immediately after a switch trial. After the break, the task block continued with the same task cued as in the switch before the break. The last trial before the break and the first trial after the break were included for purposes of calculating run length before the next switch. This created a total of 8 switch 4 trials, 8 switch 6 trials, and 8 switch 12 trials, with the remainder being repeat trials and the two start-up trials. The entire task lasted approximately 18 minutes.

Picture and word stimuli were centred 3.6 degrees of visual angle left and right of screen centre. All words measured approximately 5.6 (width) x 2.1 (height) degrees of visual angle. The pictures varied in proportions with a maximum dimensions of approximately 6.0 (width) x 4.5 (height) degrees of visual angle. The cue was presented centrally measuring approximately 1.4 x 1.4 degrees of visual angle. To create the two cues, the left or right corners of a square were rounded using Adobe Photoshop. By using only a subtle difference between the two sides of the cue, we aimed to avoid a salient visual change that would alert participants to task switches.

There were equal numbers of “yes” and “no” trials for each combination of task and switch type. At the task switch trial and the first stay trial after a task switch, correct answers for the word task and the picture task were always different. For all other stay trials, the correct response for the cued task was randomised so that it was the same as the uncued task on half of the trials and different on the other half of the trials.

Before the start of the task, participants were shown 2 trials on paper and asked which task they would do given the cue shape and what button they would press in response to the cued stimulus. Participants continued with the main task only when the experimenter was satisfied that the participant understood the instructions. No subject needed more than two repetitions of the instructions before continuing with the main task.

Scores concerned proportion error and median correct RT on switch trials. To capture the effect of time since last switch, scores were proportion error on switch 12 trials minus proportion error on switch 4 trials, and similarly RT on switch 12 minus RT on switch 4.

### Neuroradiological Assessment

MRI T1 and T2 structural scans were acquired for all patients. Lesions were traced on structural images by a neurologist, blind to the experimental results, using MRIcron (Rorden and Brett 2000) before normalising to MNI space using SPM software (Wellcome Department of Imaging Neuroscience, London, England; www.fil.ion.ucl.ac.uk) with cost-function masking to mask the lesion from the calculation of the normalization parameters (Brett et al. 2001).

### Regions of Interest

Volumes for the MD network, DMN and all other grey matter regions were constructed in the following steps. The resulting MD network (red) and DMN (blue) regions of interest (ROIs) are presented in Figure 4a. Figure 4b shows patient lesion overlap.

**Figure 4.**
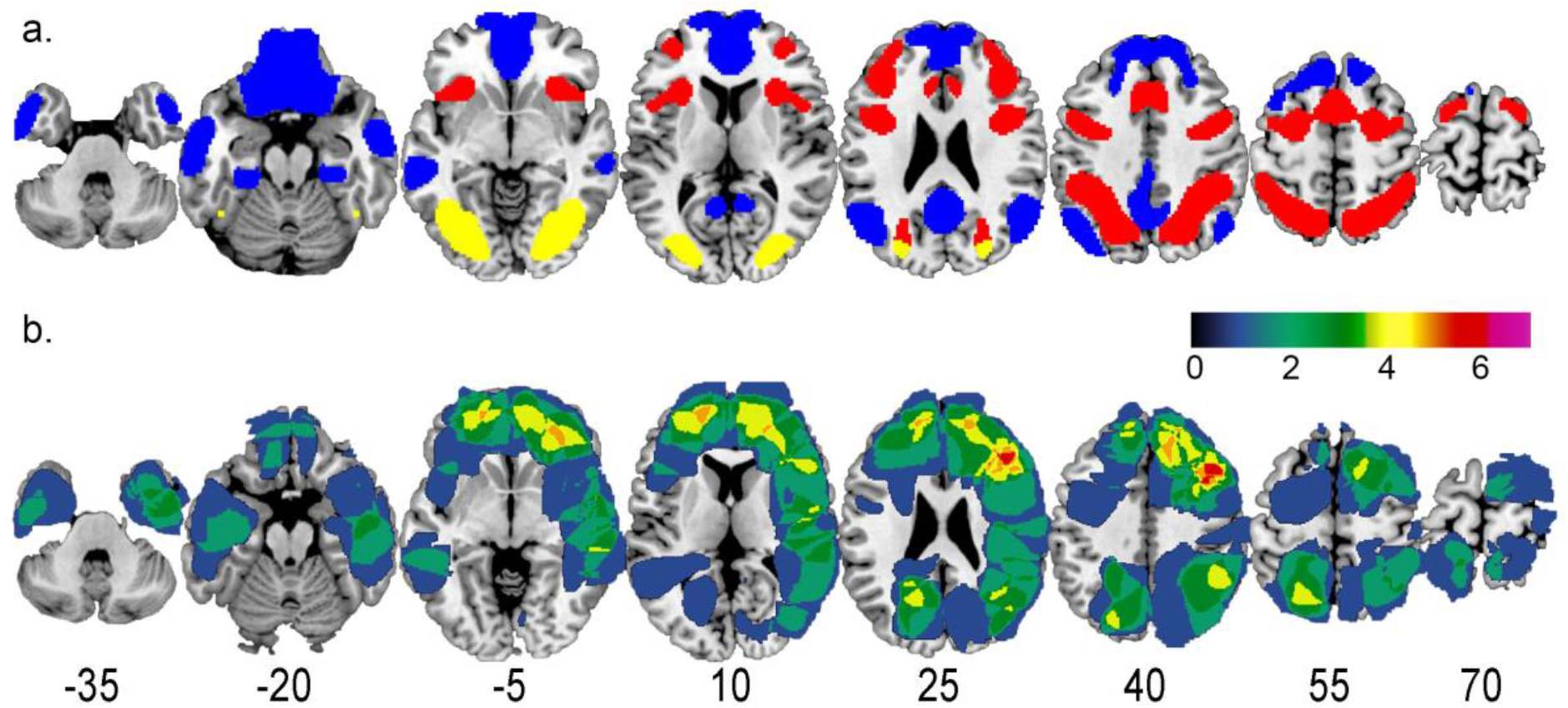
a. ROI volumes for frontoparietal MD network (red), DMN (blue) and the excluded occipital-temporal region (yellow). b. Patient lesion overlap. Colour bar shows number of patients. Numbers for each slice show z coordinate in MNI space. Apparent extension of some lesions outside the brain reflects limitations of scan quality, tracing and/or normalization.

The MD ROI was based on data from Fedorenko et al. (2013) (their figure 2), selecting only frontoparietal regions. This frontoparietal MD ROI (Figure 4a, red) included the posterior–anterior extent of the inferior frontal sulcus (pIFS, aIFS), a posterior dorsal region of lateral prefrontal cortex (pdLFC), inferior frontal junction (IFJ), anterior insula/frontal operculum (AI/FO), presupplementary motor area/dorsal anterior cingulate (preSMA/ACC), and intraparietal sulcus (IPS). A template for these regions was downloaded from http://imaging.mrc-cbu.cam.ac.uk/imaging/MDsystem. These regions were combined to create one MD volume.

The DMN ROI, presented in blue in Figure 4a, was generated using the 3 DMN networks from the liberal mask of the 17 network cortical parcellation reported in Yeo et al. (2011) (networks 15, 16 and 17). Additionally, network 10, containing temporopolar and orbital frontal regions was also included, as it contained ventromedial prefrontal cortex (vmPFC), a region often included in the DMN (Andrews-Hanna et al. 2010). These 4 networks were combined and smoothed with a 4mm FWHM Gaussian smoothing kernel and voxels with values > 0.5 after smoothing were retained.

As the MD volume AI/FO and the DMN volume vmPFC showed slight overlap, the region of overlap was removed from both ROIs.

The Other region was created using custom scripts for SPM 12 (Wellcome Department of Cognitive Neurology, London, UK). First a whole grey matter volume was created by concatenating all grey matter regions included in the AAL Atlas (Tzourio-Mazoyer et al. 2002). Then grey matter included in the MD or DMN ROIs, and all grey matter 5 mm or less from these volumes, was excluded. An additional occipital-temporal region is sometimes associated with the MD network but also strongly related to visual processing, and therefore this region (Figure 4a, yellow) was also excluded. Remaining grey matter was assigned to the Other ROI. For each patient, volumes of damage were separately measured in MD, DMN and Other ROIs.

## Results

### Differences between patients and controls

Scores for each task are shown in Table 2 (left), separately for patients and controls. One-way analyses of variance (ANOVAs) showed that patients performed significantly worse than controls on all but one measure (Table 2, middle). To assess the role of fluid intelligence, we used analysis of covariance (ANCOVA) with Culture Fair IQ as a covariate. Results are shown in Table 2 (right), along with values of Pearson’s r, derived from appropriate variance terms in the ANCOVA, reflecting the average within-group association between the variables. As predicted, though scores on two tasks (Hotel, Situations RT) were correlated with fluid intelligence, removing the effect of fluid intelligence left all significant group differences intact. Scatterplots relating each score to Culture Fair IQ are shown in Figure 5, with regression lines for each group calculated from the ANCOVA and constrained to have the same slope in the two groups.

**Table 2.** Descriptive statistics and group differences for patients and controls. The table presents mean scores and standard deviations (SD) for patients and controls in the 6 task measures, followed by group differences between patients and controls before (ANOVA) and after (ANCOVA) accounting for differences in Culture Fair IQ. Asterisks (*) represent significant effects (p<0.05). d.f. = degrees of freedom, F = F-value, P = p-value, r = within-group Pearson’s correlation with IQ.

|  | Patients |  | Controls |  | ANOVA |  |  | ANCOVA |  |  |  |
| --- | --- | --- | --- | --- | --- | --- | --- | --- | --- | --- | --- |
|  | Mean | SD | Mean | SD | d.f. | F | P | d.f. | F | P | r |
| **Culture Fair (-IQ)** | -96 | 22 | -107 | 15 | (1,60) | 5.07* | 0.028 | - | - | - | - |
| **Hotel (time deviation, sec)** | 335 | 277 | 158 | 109 | (1,60) | 10.78* | 0.002 | (1,59) | 6.53* | 0.013 | 0.34* |
| **Situations (proportion errors)** | 0.17 | 0.08 | 0.12 | 0.05 | (1,60) | 7.47* | 0.008 | (1,59) | 4.47* | 0.039 | 0.26 |
| **Situations (RT, sec)** | 10.5 | 5.69 | 6.95 | 1.66 | (1,60) | 10.86* | 0.002 | (1,59) | 6.15* | 0.016 | 0.41* |
| **Switch Time 12 – 4 (proportion errors)** | 0.08 | 0.17 | 0.02 | 0.11 | (1,56) | 2.77 | 0.101 | (1,55) | 3.29 | 0.075 | -0.12 |
| **Switch Time 12 - 4 (RT, sec)** | 0.66 | 0.91 | 0.00 | 0.36 | (1,56) | 13.27* | <0.001 | (1,55) | 11.57* | 0.001 | 0.10 |

**Figure 5.**
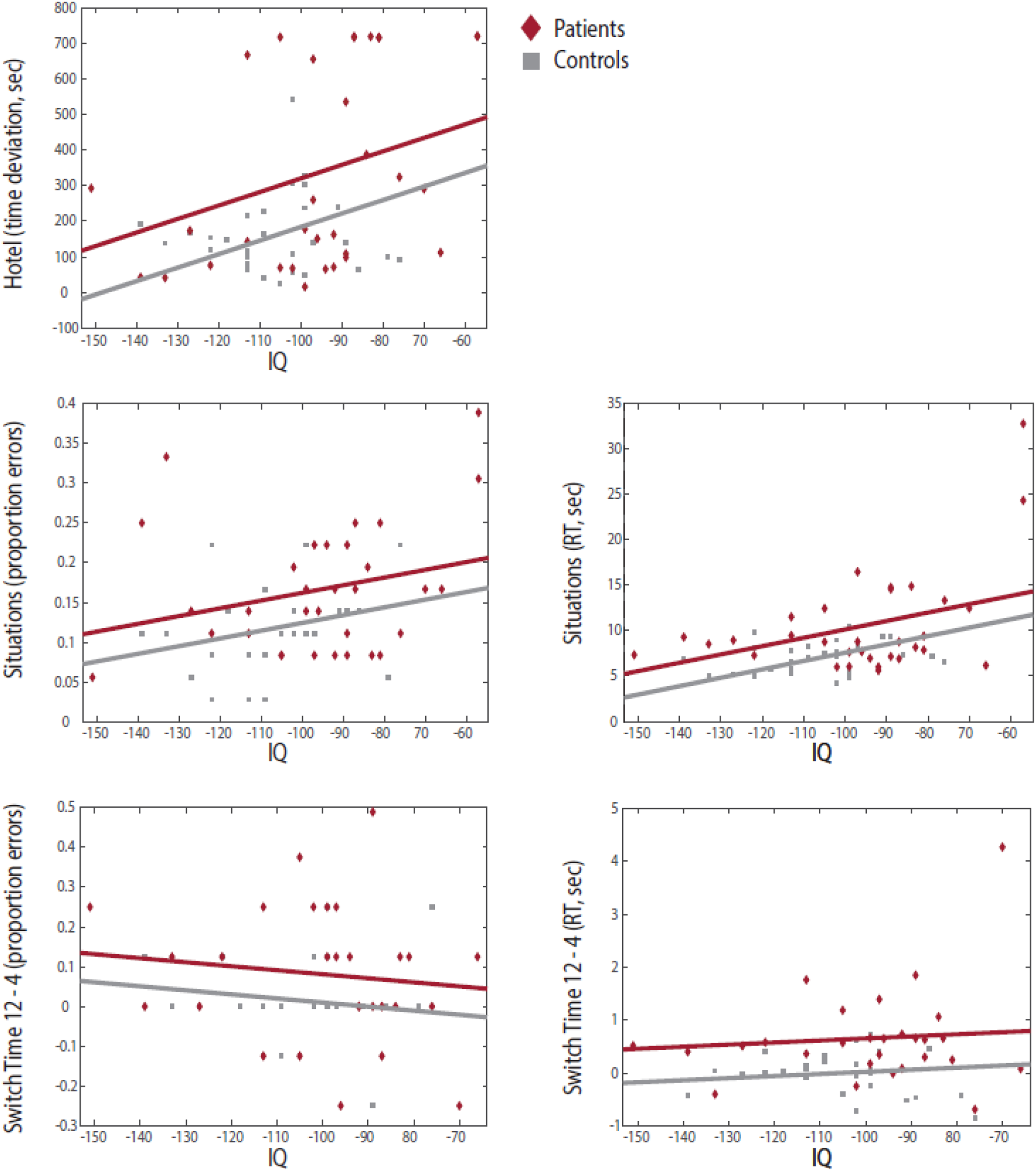
Scatterplots showing the relationship between Culture Fair IQ and the five naturalistic task measures within patients and controls separately. Regression lines reflect the average within group association of the two variables, as determined by the ANCOVAs, constrained to have the same slope across groups.

As a direct comparison with Roca et al. (2010) these analyses were re-run for the 14 patients (12 for the Switch Time task) whose lesions were restricted to the frontal lobe. Results were similar to those obtained in the full group, except that, in the ANCOVA, the significant group difference for Situations, proportion error was removed.

For Switch Time, our primary scores concerned the effects of preceding block length on switch trials (difference between switch 12 and switch 4 trials). In this task, however, we note that patients also showed substantial impairments even on stay trials (mean proportion errors 0.06 and 0.04 respectively for patients and controls, p < 0.08; median RT 2.40 and 1.57 sec, p < 0.004).

### Effects of lesion volume

The next analysis examined the relationship between behavioural scores and lesion volumes. This analysis was restricted just to the patient group. For MD, DMN and Other ROIs, the mean (standard deviation) volumes of damage were 8.39 (7.32), 8.43 (11.15) and 10.65 (7.76) ml respectively. Across patients, volumes of damage in the 3 ROIs were close to independent (maximum correlation 0.14).

Following Woolgar et al. (2010, 2018), we asked first whether Culture Fair IQ was predicted by volume of MD lesion. Scatterplots relating IQ to ROI lesion volumes are shown in Figure 6. To account for multiple comparisons (ROIs), significance threshold for correlations was set to p<0.017, one-tailed. Consistent with Woolgar et al. (2010, 2018), patient IQ was found to be significantly related to MD lesion volume (r=0.53, p<0.001). There was no relationship to DMN lesion volume (r=- 0.21, p=0.88), and only a weak association with Other lesion volume (r=0.35, p=0.03). Total lesion volume was also not significantly related to IQ (r=0.19, p=0.15).

**Figure 6.**
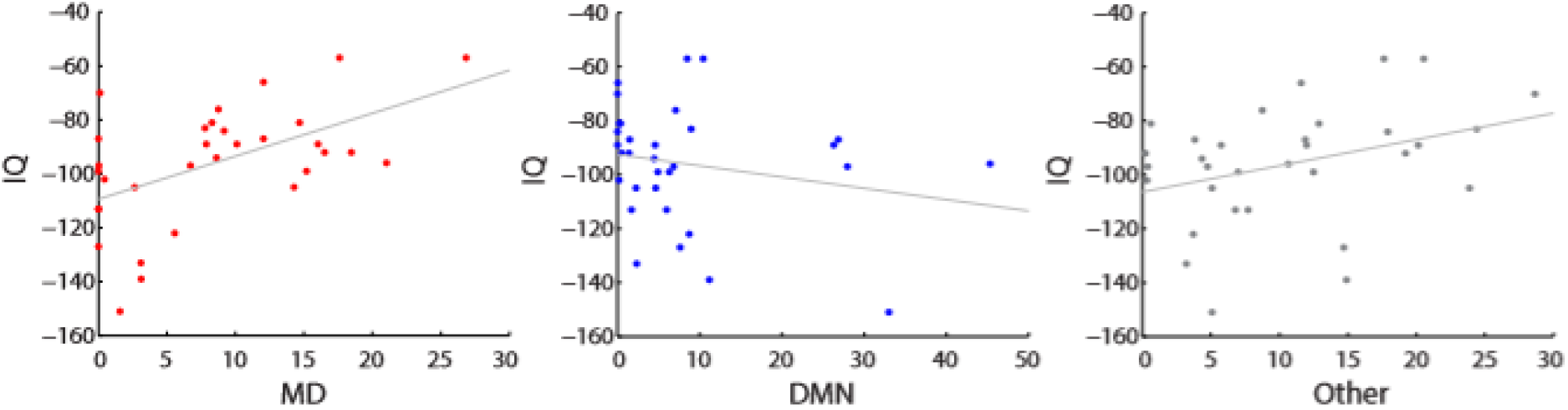
Scatterplots relating Culture Fair IQ to MD, DMN and Other lesion volumes (ml).

As 9 of the current patients were also tested in Woolgar et al. (2010), the correlation between Culture Fair IQ and MD lesion volume was re-calculated with those 9 patients excluded. Results for this independent patient group replicated the Woolgar et al. (2010) findings (r=0.39, p=0.03).

Similar analyses were then carried out for naturalistic task scores. The results are shown in Table 3. No measure was significantly related to lesion volume in either MD or DMN regions. For the RT score (switch 12 - 4) from the Switch Time task, there was a significant correlation with lesion volume in the Other ROI. The naturalistic measures were also unrelated to total lesion volume.

**Table 3.** Correlations (Pearson’s r) between lesion volume within each ROI and naturalistic task performance. Significant values (one-tailed, correcting for multiple comparisons across 3 ROIs) are shown with an asterisk (*).

|  | MD | DMN | Other |
| --- | --- | --- | --- |
| **Hotel (time deviation, sec)** | 0.23 | -0.06 | 0.11 |
| **Situations (proportion errors)** | 0.28 | -0.15 | 0.11 |
| **Situations (RT, sec)** | 0.27 | 0.09 | 0.33 |
| **Switch Time 12 – 4 (proportion errors)** | -0.15 | -0.12 | -0.22 |
| **Switch Time 12 - 4 (RT, sec)** | -0.19 | 0.05 | 0.56* |

### Between-task correlations

To test whether the naturalistic tasks were related to each other, between-task correlations were performed separately in patient and control groups. Results are shown in Table 4, with patient group correlations in the top triangle and control group correlations in the bottom triangle. With only a single exception (patient group, correlation of Situations RT with Switch Time RT difference), correlations were uniformly nonsignificant.

**Table 4.** Between task correlations (Pearson’s r) for patients (top triangle) and controls (bottom triangle). Asterisks (*) represent significant two-tailed correlations (p<0.05).

|  | **Hotel (time deviation, sec)** | **Situations (proportion errors)** | **Situations (RT, sec)** | **Switch Time 12 – 4 (proportion errors)** | **Switch Time 12 - 4 (RT, sec)** |
| --- | --- | --- | --- | --- | --- |
| **Hotel (time deviation, sec)** |  | 0.23 | 0.31 | -0.33 | -0.01 |
| **Situations (proportion errors)** | -0.08 |  | 0.35 | -0.20 | -0.24 |
| **Situations (RT, sec)** | 0.15 | 0.03 |  | 0.08 | 0.43* |
| **Switch Time 12 – 4 (proportion errors)** | 0.23 | 0.36 | -0.15 |  | -0.21 |
| Switch Time 12 - 4 (RT, sec) | 0.02 | -0.21 | -0.03 | -0.02 |  |

### Hemispheric asymmetry

To check for differential involvement of left and right hemispheres in our naturalistic tasks, we compared performance in patients with left (n = 11) and right (n = 18) hemisphere lesions (see Table 2). For naturalistic tasks there was no evidence of hemispheric differences (maximum t = 0.99), contrasting with worse performance for the right hemisphere group in the Culture Fair, t = 2.39, p < .02.

### Residual patient impairment

In a final examination of lesion effects, we calculated average performance across naturalistic tasks, after accounting for effects of fluid intelligence. For each patient, standardized residual scores, calculated from the above ANCOVAs, were derived for all 5 naturalistic-task scores and then averaged. Signs were set such that high scores reflected poorer performance than predicted from Culture Fair IQ. For the four patients who had not completed the Switch Time task, the average residual score was generated from Hotel and Situations scores. After correcting for multiple comparisons, the average residual was not correlated with volume of damage in any ROI, with the strongest correlation for Other (r=0.31). Figure 7a shows the lesions of the 6 patients with the highest average residual scores, reflecting naturalistic task performance worse than predicted from fluid intelligence. Rather than implicating a specific brain region, the figure illustrates the diversity of lesions associated with naturalistic task impairment.

**Figure 7.**
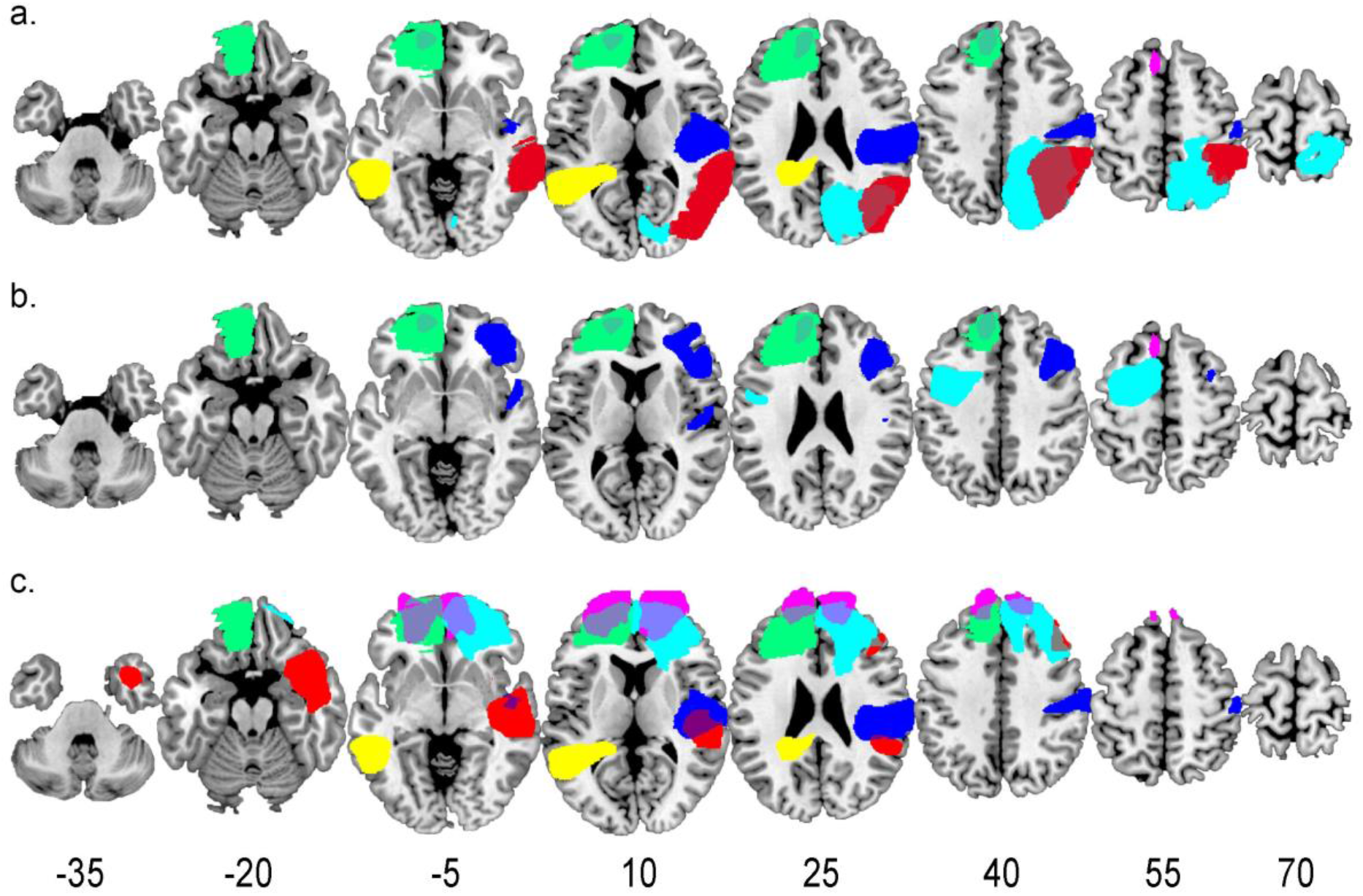
Lesion overlap for the patients with the greatest residual impairment after accounting for differences in Culture Fair IQ. a. Shows the lesions of the 6 most impaired patients from the whole sample. b. Shows the lesions of the 4 most impaired patients when the patient sample was restricted the patients with only lesions to the frontal lobe. c. Shows the lesions of the 6 most impaired patients on Switch Time only. For each subplot, each colour denotes a different patient. Numbers for each slice show z coordinate in MNI space.

In order to compare with Roca et al. (2010, 2011) we recalculated patient residuals after restricting the patient sample to patients with frontal lesions only. Figure 7b shows the lesions of the 4 frontal patients with the highest average residual scores. Unlike Roca et al. (2010), residual impairment after accounting for IQ was not restricted to the anterior PFC but spread across the frontal lobe. It should be noted however, that this analysis was done on a small sample of 14 patients only.

As the Switch Time RT score was significantly related to damage in the Other ROI, we conducted a final similar analysis to search for a common lesion location among patients with the greatest deficits on this score only. The analysis was the same as for average residual scores (Figure 7a), but this time using only the Switch Time residual. Again the results showed scattered brain lesions among the 6 patients with the greatest deficit (Figure 7c).

## Discussion

In previous work, we have found that “executive” tasks vary widely in the degree to which deficits are explained by fluid intelligence (Roca et al. 2010, 2012; Woolgar et al. 2010). For several classical tests, such as Wisconsin card sorting, impairments in diverse patient groups are eliminated when fluid intelligence is partialled out. The Hotel task of Manly et al. (2002), however – based on the 6-element task of Shallice and Burgess (1991) – shows a very different result. For this task there is only a weak correlation with fluid intelligence, and patient impairments are not removed by partialling out this effect. These results match the long-held belief that complex, naturalistic tasks can reflect aspects of cognitive deficit that are missed in classical tests.

In the present work, we confirmed these findings for a new version of the Hotel task. Again, deficits in a diverse group of patients with focal brain lesions were not accounted for by fluid intelligence. Results were similar in a second test of understanding and decision-making in complex, life-like scenarios. In a third test, we attempted to isolate one critical factor in the Hotel task and potentially other real-world situations – the need to break out from a lengthy period of immersion in a single task. While task switching in the Hotel is spontaneous, with no external cue, we used an explicit switch cue but varied the length of the previous task block. Here too we observed deficits in the patient group that were not explained by fluid intelligence. Though fluid intelligence may account for deficits in many “executive” tasks, our results extend the list of more naturalistic tasks for which this is not true.

Also replicating previous work, we found that deficits in fluid intelligence were predicted by the extent of lesions to the frontoparietal MD network. MD lesions, however, were not strong predictors of deficits in the naturalistic tasks. While MD lesions may explain many aspects of classical executive deficit, again these data suggest a different explanation for naturalistic deficits.

As a second potential predictor of naturalistic deficits, we considered lesions to the DMN. Influential accounts link the DMN to representation of broad cognitive contexts, including spatial, temporal and social aspects (e.g. Ranganath and Ritchie, 2012, Andrews-Hanna et al., 2010). Plausibly, broad contextual representations will be especially important in temporally extended, open-ended behaviour. As we found for the MD network, however, naturalistic impairments were not strongly predicted by DMN lesions.

Instead, we found that naturalistic deficits could arise from lesions scattered through multiple regions of the brain, in left or right hemispheres, in frontal, parietal or occipito-temporal cortex, and including or not sections of MD and DMN networks. Though performance was not significantly associated with either MD or DMN lesions, our interpretation is not that these networks make no contribution to naturalistic behaviour. Rather, no one network is strongly predictive of deficit simply because deficits can arise from lesions of many different kinds.

Our results for Hotel and Situations are firmly in line with the finding that complex, naturalistic tasks capture cognitive deficits missed in many executive tests, but suggest that these tests will be poor at isolating any specific cognitive process. In line with the arguments of Burgess et al. (2000), complex, real-life decision making rests on many processes, likely dependent on many different cortical regions. The particular processes that are critical may vary between tasks, matching our finding of generally nonsignificant correlations between them. Perhaps not surprisingly, complex, real-life decision-making may depend on much of the cortex, making these tests highly sensitive to brain lesions, but not highly diagnostic of any specific cognitive deficit.

For Switch Time this argument is less clear, since the test is ostensibly much more focused in its cognitive demands. Results however were similar, with deficits not explained by fluid intelligence, and unrelated to volume of damage in either MD or DMN regions. For this test there was a significant correlation with damage to the residual, Other ROI, but given widely scattered lesions in the most impaired patients (Figure 7c), along with differences in total size of MD, DMN and Other regions this result should be interpreted with caution. Further work would be needed to analyse deficits in this task. By design, the cue indicating a change of task had low visual salience, perhaps requiring participants to maintain a sustained attentional set for its possible occurrence. The duration of a block of 12 trials depended on individual RTs, with typical durations around 40 s for controls, but appreciably greater for patients given slow responses even on stay trials. The data suggest that escape from a period of immersion in one task indeed calls on cortical functions unlike those captured in fluid intelligence and, by extension, other classical executive tests. Again, however, deficits may have mixed causes, not simply related to specific lesion locations.

In principle, the source of a deficit in any one task and patient might be clarified by extensive neuropsychological testing. Given the low correlations between our naturalistic tasks, however, it seems unlikely that any small set of cognitive processes will be broadly predictive of “naturalistic-task” deficits. Instead, we suggest that each such task has its own, rich cognitive profile, likely impaired for different reasons in different patients.

On this interpretation, naturalistic tasks do not identify specific control deficits required only in complex, open-ended behaviour. Instead, their sensitivity to brain damage arises simply through the many cognitive processes that must be combined, bringing many opportunities for performance impairment. In line with the original work of Shallice and Burgess (1991), such tests may be especially useful as an indication of difficulties that patients may face in return to everyday life. To predict such difficulties, clinicians should use not only measures that target specific cognitive abilities, such as classical executive tasks which are associated with lesion to the MD system (Roca et al. 2010), but also tasks – like the ones described here - that tackle more complex, naturalistic scenarios. This way, we can provide a more comprehensive cognitive assessment which could better reflect real life deficits. Regarding neuropsychological rehabilitation, the findings shed light on the complicated task that neuropsychologists face when dealing with brain lesions. As clinical experience shows, difficulties in returning to everyday life are not easily predicted from damage to specific brain regions or networks, reflecting the complexity of cortical involvement in everyday behavioural management and decisions.

## Acknowledgments

This work was funded by the UK Medical Research Council (intramural program SUAG/045.G101400) and the British Academy (grant SG160590).

## Supplementary Material

### Situations Task – Full Question Set

* = correct answer, ^ = second best answer.

Gabrielle has recently changed jobs and she’
ss going through a financial struggle. Her sister Amy’s birthday was coming up so she had been walking to work in order to save the bus fare to get her something pretty. She finally saved up a little and bought Amy a bracelet. The day of the birthday Gabrielle was at her sister’s house with some of Amy’s friends. After Amy opened all the presents, Gabrielle decided to start doing the dishes in order to help Amy out. She then heard Amy saying to her friends that she did not like the bracelet, because it looked cheap and tacky. After a few days the bills finally arrived, and Gabrielle noticed that she had spent more money on the bracelet than she should have. Now she didn’t have enough money to pay her bills.

#### Executive question

In order to pay her bills Gabrielle should:

1. Pawn her jewellery^
2. Rob a bank
3. Work out which bills can be put off until her next pay check*

#### Feelings question

When Gabrielle heard Amy she felt:

1. Hurt*
2. Perplexed^
3. Inspired

#### Social question

She should:

1. Trash Amy’
ss room
2. Wait until everybody has left and tell Amy how she felt*
3. Get angry at Amy and not speak to her for a while^

Josephine and David are getting married in November. They have agreed to a simple wedding in David’
ss parents’ garden, with a few close friends and relatives, because they are saving to buy a house in Canterbury. The garden is small, but they have planned to fit in 4 tables for the guests they have invited. They are eating out at a nice restaurant, while discussing the final arrangements of the wedding. Suddenly, Josephine receives an email from one of her college friends saying they heard about the wedding and 5 of them have booked flights to participate in it.

#### Social question

For the wedding, David should wear:

1. A dress
2. A dinner jacket^
3. A suit*

#### Feelings question

About this situation Josephine felt:

1. Nervous^
2. Concerned*
3. Rejected

#### Executive question

Regarding the space issue, Josephine should:

1. Try to fit in an extra table*
2. Cancel the wedding
3. Move the meal to a nearby park^

Nick was on holiday at an all-inclusive resort in Playa del Carmen, México. One beautiful and warm morning he decided to go for a walk to get to know the place a little better. The water was so clear, he could see both his feet and the fish swimming around them. As he walked by another resort, he happened to run across his boss’
ss boss, Colin. Colin recognized Nick, and asked him to join Colin and his young wife for dinner that night at one of the 6 restaurants in Colin’s hotel.

#### Feelings question

Regarding Colin’s invitation Nick felt:

1. Excited*
2. Worried^
3. Empty

#### Executive question

Nick did not remember the name of the restaurant. He should:

1. Go to MacDonald’s alone
2. Call the hotel and ask to speak to Colin’s room to ask him the name of the restaurant*
3. Go to every restaurant in the hotel and try to see if Colin is there^

#### Social question

When he meets Colin’s wife, Nick should be:

1. Friendly*
2. Flirtatious
3. Very Formal^

Michael’
ss son Trevor had been asking him to teach him how to drive for a very long time now. When Trevor turned 19 and got his first job Michael thought it was time and started teaching him with his brand new Toyota Corolla. Trevor was a very good driver, respectful and prudent. On their way back from Trevor’s driving test, Trevor asked Michael if he could drive the last block. Michael agreed and changed places with him. As soon as Trevor started driving a dog appeared, he tried to dodge it and crashed into his neighbor’s car. Michael could not immediately afford the cost of repairs, and had been planning to drive friends to Edinburgh in two weeks’ time.

#### Executive question

To deal with costs Michael should:

1. Borrow money for the repairs on his credit card*
2. Sell his car
3. Wait a few months until he can save enough money^

#### Social question

Michael should tell Trevor to:

1. Ring the neighbor’s bell, tell him what happened, apologize and offer to pay for the damages*
2. Leave a note with the insurer’s details but not mention he was the driver^
3. Escape the scene

#### Feelings question

About the accident Trevor felt:

1. Loved
2. Anxious^
3. Ashamed*

Sonia has just started at a new job. She was working by herself as a party planner but got fed up and decided to try something different. As she has majored in marketing she applied for a position in a company and luckily got the job. She started on Monday and on Friday she was invited to a party at the house of one of her co-workers. The gathering was on the other side of town and it took her forever to get there. As she was arriving she looked for her phone to check the exact apartment when she notice that it wasn’
st there. She had left it charging in the bathroom! She was able to remember the address but not the apartment number.

#### Feelings question

Sonia felt:

1. Useless^
2. Angry with herself*
3. Jubilant

#### Executive question

Sonia should:

1. Ring one of the apartments and ask if they know her co-worker’
ss apartment number^
2. Go back home
3. Look around to see if someone else is arriving*

#### Social question

For the party, she should bring:

1. A nice bottle of wine*
2. A can of beer^
3. A Christmas pudding

Samantha has two sons, Jack aged 5 and Thomas aged 2. That afternoon she picked up Jack from school and took him to soccer practice. She took Thomas with her as well in order to take him to the park for a while. She was returning from the park on her way to pick up Jack from the football field, when she drove past a meat market and decided to stop to buy some meat for dinner. As soon as she got out, the car locked behind her with Thomas still inside. She tried everything she could think of but couldn’
st open the doors and Thomas started crying. At the same time she knew Jack would be waiting on his own.

#### Feelings question

When the door locked Samantha felt:

1. Cheerful
2. Enraged^
3. Desperate*

#### Social question

To make arrangements for Jack she should:

1. Call her husband and ask him to pick Jack up*
2. Call the school and ask if a teacher will look after him^
3. Think it’s time for him to grow up anyway

#### Executive question

Samantha should:

1. Look for something to break a window^
2. Buy the meat and deal with this later
3. Find the number of a local locksmith*

Tim had been planning a trip to Salzburg for 6 months. He read a lot of reviews about where to go and what to eat. He flew on Wednesday to Salzburg, and arrived really early. Since the check in at the hotel started at 12.00am he decided to grab a bite in the meantime. He finally arrived at the hotel he had booked at 11.55am feeling very tired from walking around. Upon his arrival he was told at the front desk that they had no reservation under his name and no rooms available because of a convention happening that week. They offered to find him a room in another hotel, but this was much more expensive and completely beyond his budget.

#### Social question

Tim should:

1. Ask to speak to the manager and try to solve the situation*
2. Punch the receptionist
3. Shout at the receptionist and give the hotel a bad review^

#### Executive question

If there really is no room, Tim should:

1. Try to find another hotel at the right price*
2. Stay at the new hotel he was offered but cut his visit short^
3. Fly back home

#### Feelings question

Tim felt

1. Confused
2. Bored^
3. Annoyed*

Stella has a very good relationship with her mother-in-law, Frances. Frances really helps Stella with the kids, but she is also very talkative. Stella is at her work at the bank and she has a very busy day as a Director is visiting from France. Frances has been wanting to talk with her for days about what decorations she can buy for one of the children’s birthday parties, which is going to take place tomorrow. Stella is in the middle of an urgent report when she sees her mother-in-law calling her cellphone. She wants to take the call because the birthday is tomorrow but has no time to discuss the decoration right now. If she does not answer Frances, she will probably end up having to buy everything herself and having to bake the cake herself as well.

#### Feelings question

When Stella sees the call she feels:

1. Torn*
2. Proud
3. Uneasy^

#### Social question

Stella should:

1. Pick up the phone and tell her mother-in-law she won’t be able to talk right now*
2. Ignore her call and deal with her when she gets out of work^
3. Quit her job

#### Executive question

If she can’t get Frances’ help, Stella should:

1. Have no decorations for the birthday party^
2. Cancel the birthday party
3. Buy the decorations and a prepared cake from a shop*

Greg and Scarlett had been planning a trip to Thailand for a long while. They thought and took care of every little detail: flights, hotels, tours, transportation, they even arranged with Greg’
ss father to take care of Rocko, their dog, while they were away. A day before their trip, Greg’s father called them and told them he had decided to go on a last minute trip and wouldn’t be able to take care of Rocko. They called a few relatives and friends about looking after the dog, but without success.

#### Social question

Responding to his father, Greg should:

1. Get mad at his dad^
2. See if there is any way he can postpone his trip*
3. Break down in tears

#### Feelings question

Talking to his friends, Greg felt:

1. Frustrated*
2. Disconcerted^
3. Fulfilled

#### Executive question

To deal with Rocko they should:

1. Postpone the trip^
2. Put Rocko in a kennel*
3. Give the dog away

Benjamin’s wife is having a surprise birthday party for her sister. She has asked Benjamin to pick up the cake and present she ordered, because the stores are really close to his office. She advised him to be on time because the stores close at 5pm. Benjamin gets caught up with work and forgets about his errand. On his way out of the office at 5.30pm he realizes he has not bought what he was asked for. Instead he buys pudding in a nearby store and arrives to the birthday party late, with no cake or present.

#### Executive question

Benjamin should have:

1. Used his lunch break to buy all the things in advance*
2. Skipped the party and gone golfing
3. Set an alarm for 4.50 to go to the shops^

#### Social question

When he gets to the party Benjamin should:

1. Apologize to his wife and sister-in-law for what has happened*
2. Have a nap in his car and then go in
3. Say they should have fun anyway^

#### Feelings question

Benjamin’s wife feels:

1. Ashamed^
2. Safe
3. Angry*

Charlie and Peter went to see a movie. As Peter had an early dinner they caught the last show. As they were getting out of the cinema, Charlie asked Peter if he could wait while he went to the restroom. Peter was very tired so he declined, said goodbye and went his own way. Charlie went into the bathroom and when he tried to get out he realized the door was locked. He shouted, but nobody responded. It was very late and there were few people left at the cinema.

#### Executive question

To resolve this situation he should:

1. Sit and wait until someone appears^
2. Continue shouting and knocking on the door*
3. Smash the window and try to jump down into the street

#### Feelings question

Charlie felt:

1. Perplexed^
2. Frustrated*
3. Alive

#### Social question

When Charlie finally escapes he should:

1. Write a letter to the cinema explaining what happened and ask for an apology*
2. Never go to that cinema again^
3. Set the cinema on fire

George is on his way to work, where he has an important meeting with a potential client. He lives an hour away from his work and really enjoys driving there every day, because he drives through really beautiful places. As he has almost reached the city center, he realizes he has forgotten his laptop with the presentation at home. If he goes back to get it he will be very late for the meeting. At that moment his wife calls him and tells him she noticed he had left behind his laptop. She is already in the car bringing it to him, but she will get there probably half an hour after the meeting has started.

#### Social question

Because he doesn’t have his presentation with him George should:

1. Explain what happened and ask to move the meeting half an hour*
2. Make up an excuse (e.g. car problems) and wait until his wife arrives^
3. Buy the client a present

#### Executive question

To make sure this never happens again, George should:

1. Always leave his laptop with his car keys*
2. Move house so he is closer to work
3. Always arrive for work very early^

#### Feelings question

Speaking to his wife, George feels:

1. Delighted^
2. Grateful*
3. Sad

## References

Addis DR, Pan L, Vu MA, Laiser N, Schacter DL. 2009. Constructive episodic simulation of the future and the past: Distinct subsystems of a core brain network mediate imagining and remembering. Neuropsychologia. 47:2222–2238.

Addis DR, Wong AT, Schacter DL. 2007. Remembering the past and imagining the future: Common and distinct neural substrates during event construction and elaboration. Neuropsychologia. 45:1363–1377.

Amodio DM, Frith CD. 2006. Meeting of minds: The medial frontal cortex and social cognition. Nat Rev Neurosci. 7:268–277.

Andrews-Hanna JR, Reidler JS, Sepulcre J, Poulin R, Buckner RL. 2010. Functional-Anatomic Fractionation of the Brain’s Default Network. Neuron. 65:550–562.

Barbey AK, Colom R, Solomon J, Krueger F, Forbes C, Grafman J. 2012. An integrative architecture for general intelligence and executive function revealed by lesion mapping. Brain. 135:1154–1164.

Benton AL. 1968. Differential behavioral effects in frontal lobe disease. Neuropsychologia. 6:53–60.

Brainard DH. 1997. The Psychophysics Toolbox. Spat Vis. 10:433–436.

Brett M, Leff AP, Rorden C, Ashburner J. 2001. Spatial normalization of brain images with focal lesions using cost function masking. Neuroimage. 14:486–500.

Burgess PW, Veitch E, De Lacy Costello A, Shallice T. 2000. The cognitive and neuroanatomical correlates of multitasking. Neuropsychologia. 38:848–863.

Christoff K, Gordon AM, Smallwood J, Smith R, Schooler JW. 2009. Experience sampling during fMRI reveals default network and executive system contributions to mind wandering. Proc Natl Acad Sci U S A. 106:8719–8724.

Crittenden BM, Mitchell DJ, Duncan J. 2015. Recruitment of the default mode network during a demanding act of executive control. Elife. 4:e06481.

D’Argembeau A, Collette F, Van Der Linden M, Laureys S, Del Fiore G, Degueldre C, Luxen A, Salmon E. 2005. Self-referential reflective activity and its relationship with rest: A PET study. Neuroimage. 25:616–624.

Fedorenko E, Duncan J, Kanwisher N. 2013. Broad domain generality in focal regions of frontal and parietal cortex. Proc Natl Acad Sci U S A. 110:16616–16621.

Frith U, Frith CD. 2003. Development and neurophysiology of mentalizing. Philos Trans R Soc B Biol Sci. 358:459–473.

Gaffan D. 1994. Scene-Specific Memory for Objects: A Model of Episodic Memory Impairment in Monkeys with Fornix Transection. J Cogn Neurosci. 6:305–320.

Gläscher J, Rudrauf D, Colom R, Paul LK, Tranel D, Damasio H, Adolphs R. 2010. Distributed neural system for general intelligence revealed by lesion mapping. Proc Natl Acad Sci. 107:4705LP–4709.

Hassabis D, Maguire EA. 2007. Deconstructing episodic memory with construction. Trends Cogn Sci. 11:299–306.

Institute for Personality and Ability Testing. 1973..

Manly T, Hawkins K, Evans J, Woldt K, Robertson IH. 2002. Rehabilitation of executive function: Facilitation of effective goal management on complex tasks using periodic auditory alerts. Neuropsychologia. 40:271–281.

Mason MF, Norton MI, Van Horn JD, Wegner DM, Grafton ST, Macrae CN. 2007. Wandering minds: The default network and stimulus-independent thought. Science (80-). 315:393–395.

Milner B. 1963. Effects of Different Brain Lesions on Card Sorting: The Role of the Frontal Lobes. Arch Neurol. 9:90–100.

Ranganath C, Ritchey M. 2012. Two cortical systems for memory-guided behaviour. Nat Rev Neurosci. 13:713–726.

Reitan RM. 1955. The relation of the Trail Making Test to organic brain damage. J Consult Psychol. 19:393–394.

Roca M, Manes F, Cetkovich M, Bruno D, Ibáñez A, Torralva T, Duncan J. 2014. The relationship between executive functions and fluid intelligence in schizophrenia. Front Behav Neurosci. 8.

Roca M, Manes F, Chade A, Gleichgerrcht E, Gershanik O, Arévalo GG, Torralva T, Duncan J. 2012. The relationship between executive functions and fluid intelligence in Parkinson’s disease. Psychol Med. 42:2445–2452.

Roca M, Manes F, Gleichgerrcht E, Watson P, Ibáñez A, Thompson R, Torralva T, Duncan J. 2013. Intelligence and executive functions in frontotemporal dementia. Neuropsychologia. 51:725–730.

Roca M, Parr A, Thompson R, Woolgar A, Torralva T, Antoun N, Manes F, Duncan J. 2010. Executive function and fluid intelligence after frontal lobe lesions. Brain. 133:234–247.

Roca M, Torralva T, Gleichgerrcht E, Woolgar A, Thompson R, Duncan J, Manes F. 2011. The role of Area 10 (BA10) in human multitasking and in social cognition: A lesion study. Neuropsychologia. 49:3525–3531.

Rogers RD, Monsell S. 1995. Costs of a Predictable Switch Between Simple Cognitive Tasks. J Exp Psychol Gen. 124:207–231.

Rorden C, Brett M. 2000. Stereotaxic display of brain lesions. Behav Neurol. 12:191–200.

Shallice T, Burgess PW. 1991. Deficits in strategy application following frontal lobe damage in man. Brain. 114:727–741.

Shulman GL, Fiez JA, Corbetta M, Buckner RL, Miezin FM, Raichle ME, Petersen SE. 1997. Common blood flow changes across visual tasks: II. Decreases in cerebral cortex. J Cogn Neurosci. 5:648–663.

Smith V, Mitchell DJ, Duncan J. 2018. Role of the Default Mode Network in Cognitive Transitions. Cereb Cortex. 28:3685–3696.

Torralva T, Roca M, Gleichgerrcht E, Bekinschtein T, Manes F. 2009. A neuropsychological battery to detect specific executive and social cognitive impairments in early frontotemporal dementia. Brain. 132:1299–1309.

Tzourio-Mazoyer N, Landeau B, Papathanassiou D, Crivello F, Etard O, Delcroix N, Mazoyer B, Joliot M. 2002. Automated anatomical labeling of activations in SPM using a macroscopic anatomical parcellation of the MNI MRI single-subject brain. Neuroimage. 15:273–289.

Woolgar A, Duncan J, Manes F, Fedorenko E. 2018. Fluid intelligence is supported by the multiple-demand system not the language system. Nat Hum Behav. 2:200–204.

Woolgar A, Parr A, Cusack R, Thompson R, Nimmo-Smith I, Torralva T, Roca M, Antoun N, Manes F, Duncan J. 2010. Fluid intelligence loss linked to restricted regions of damage within frontal and parietal cortex. Proc Natl Acad Sci U S A. 107:14899–14902.

Yeo BT, Krienen FM, Sepulcre J, Sabuncu MR, Lashkari D, Hollinshead M, Roffman JL, Smoller JW, Zöllei L, Polimeni JR, Fisch B, Liu H, Buckner RL. 2011. The organization of the human cerebral cortex estimated by intrinsic functional connectivity. J Neurophysiol. 106:1125–1165.

